# Heterogeneous Graph Contrastive Learning for Drug-Gene-Disease Motif Prediction

**DOI:** 10.64898/2026.09.21.752883

**Authors:** Hongsheng Xie, Yuan Gao, Shanlin Ke, Can Chen

## Abstract

Drug repurposing and target discovery offer critical strategies for advancing therapeutic development by uncovering the potential biological pathways and novel associations among drugs, genes, and diseases. However, experimental discovery remains expensive and time-consuming, which limits the scalability of large-scale studies. In addition, existing computational approaches often struggle to effectively integrate heterogeneous biomedical data, capture the complex higher-order topological signatures of biological interactomes, and generalize to unseen entities. Here, we present HANAMI (Heterogeneous grAph coNtrastive leArning for drug-gene-disease Motif predIction), a multi-view deep graph learning framework designed to model complex interactions among drugs, genes, and diseases. HANAMI integrates diverse heterogeneous biomedical knowledge, including chemical structures, genomic sequences, and clinical phenotypes, and leverages relation-aware topology encoding, structure-aware aggregation, and contrastive learning to enable accurate motif prediction with biological context from the network. Systematic evaluation on benchmark datasets shows that HANAMI achieves up to 6% improvements over existing state-of-the-art methods in predicting drug-gene-disease motifs. The framework further demonstrates strong inductive generalization, maintaining an ∼18% performance advantage in zero-shot settings involving previously unseen entities. Beyond predictive performance, HANAMI effectively prioritizes drug-disease relationships investigated in Phase II or III trials while identifying candidate genes that suggest plausible mechanistic links. Together, HANAMI provides a computational framework for interpreting complex biomedical interactions, offering a scalable foundation to accelerate drug repurposing and therapeutic innovation.

## Introduction

Many complex conditions, particularly neurodegenerative and rare diseases, still lack effective treatments [1, 2]. Addressing this unmet clinical need requires systematic characterization of interactions among drugs, genes, and diseases to better understand biological systems and advance drug repurposing and therapeutic development [3–9]. However, traditional drug discovery remains costly and time-consuming, leading to growing research interests in drug repurposing defined as the identification of new uses for existing compounds [10–14]. The effectiveness of this approach depends on a comprehensive understanding of the multi-scale relationships among drugs, genes, and diseases. Drugs act on genes to modulate biological processes underlying disease states, and mapping these tripartite interactions can reveal how pharmacological interventions influence pathways to alter disease phenotypes [15, 16]. Developing computational frameworks to resolve higher-order interaction patterns is therefore essential for drug repurposing and drug target discovery, particularly as wet-lab characterization of such interactions is often constrained by resource limitations. These frameworks enable identification of functional motifs and provide a scalable strategy for exploring previously uncharacterized regions of biomedical networks, supporting repurposing and therapeutic development.

Earlier methodologies for modeling tripartite interactions largely rely on mathematical optimization and ensemble learning to predict associations within heterogeneous graphs [17, 18]. One common approach is random forest classifiers [19], which utilize feature matrices derived from topological properties to capture nonlinear relationships in high-dimensional biological networks. Complementary strategies, such as collective matrix factorization [17], decompose multi-modal data into shared latent factors to resolve the underlying dependencies among drugs, genes, and diseases. In addition, network-based diffusion strategies, including random walk with restart [20, 21], spread information across the interactome to identify potential therapeutic indications based on network distance. More recent work has incorporated multi-view learning and genomic topology to account for the spatial organization of the interactome [22, 23]. However, these supervised and optimization-based approaches remain limited in their ability to capture complex, higher-order interaction motifs in biomedical networks and often rely heavily on manual feature engineering.

Deep graph learning methods have gained prominence for their ability to synthesize node-level attributes with the underlying network topology in biomedical prediction tasks [24–26]. Initial neural network architectures for predicting associations often utilize static embedding techniques to capture local structural information [27]. For instance, N2V-MLP [28] processes pre-computed Node2Vec embeddings through a multi-layer perceptron (MLP), but treating nodes as independent entities often fails to preserve the relational context required for triplet prediction. Subsequent methodologies incorporate specialized fusion architectures to integrate diverse data streams, such as TriNet [29] which employs parallel network branches to synthesize heterogeneous biological information. More recently, TriMoGCL [30], a state-of-the-art framework that emphasizes higher-order pattern recognition by leveraging contrastive learning [31, 32] combined with biomedical knowledge to address motif imbalance and improve predictive performance. Nevertheless, TriMoGCL is still limited by the scope of biomedical knowledge it incorporates and its ability to fully capture broader relationships between motifs and the global network structure. In particular, existing approaches do not generalize well to cases involving unseen entities. Without sufficient inductive capacity to infer relationships among multiple novel entities simultaneously, their applicability to newly discovered compounds is constrained.

To address these limitations, we propose HANAMI (Heterogeneous grAph coNtrastive leArning for drug-gene-disease Motif predIction), a multi-view deep graph learning architecture designed to identify triplet motifs within drug-gene-disease networks. HANAMI integrates multi-modal embeddings from pre-trained language models and domain-specific encoders to synthesize chemical, genomic, and clinical context, and employs a topology-aware feature refinement and structure-aware aggregation to capture both global and local relationship dynamics. We further introduce a contrastive learning objective that leverages multiple pairs of examples to improve the model’s ability to distinguish subtle structural variations and handle biological data gaps. Evaluations on the multi-scale interactome (MS) [33] and drug repurposing knowledge graph (DRKG) [34] datasets demonstrate that HANAMI consistently outperforms state-of-the-art approaches in both transductive and inductive settings, allowing for the prediction of interactions for biomedical entities not encountered during training. Beyond predictive performance, evaluation on a cohort defined using Phase II or III trial records from ClinicalTrials.gov shows that HANAMI prioritizes drug and disease relations absent from the network together with the shared genes that connect them. By identifying specific drug-gene-disease interaction patterns, the framework clarifies the pathways through which drugs interact with genes and disease-specific components, providing a foundation for drug repurposing and therapeutic discovery.

## Results

### HANAMI Architecture

The HANAMI framework is a specialized graph neural network designed to classify seven distinct structural motifs, consisting of complex triplet patterns formed by the interactions among drug, gene, and disease entities within biomedical networks. As shown in Fig. 1, the workflow contains three main phases: feature initialization, refinement, and consolidation & prediction. In the first phase, high-dimensional embeddings are generated through specific encoders, including ChemBERTa-3 [35] and MPNN (message passing neural network) [36] for drugs, Enformer [37] and Borzoi [38] for genes, and BioBERT [39] and ClinicalBERT [40]. ChemBERTa-3 reads a drug’s chemical structure as a text sequence, while the MPNN reads the same structure as a network of atoms and bonds. Together, they provide complementary views of the molecule. Enformer and Borzoi extract regulatory information from DNA sequences, whereas BioBERT and ClinicalBERT extract disease information from biomedical and clinical text. These embeddings are then subsequently integrated within a heterogeneous network using a GraphSAGE-based message passing architecture [41] that combines information about each drug, gene, or disease with information from directly connected entities. To enhance motif distinction and mitigate network noise, a contrastive learning objective [42] helps HANAMI recognize related views of the same motif and distinguish them from different interaction patterns. Finally, the model undergoes a structure-aware feature consolidation process combining information from the drug, gene, and disease as a group with separate comparisons between each pair to synthesize comprehensive triplet representations. These representations are fused and passed to a classification layer for motif prediction. Detailed descriptions of these components are provided in Methods.

**Fig. 1.**
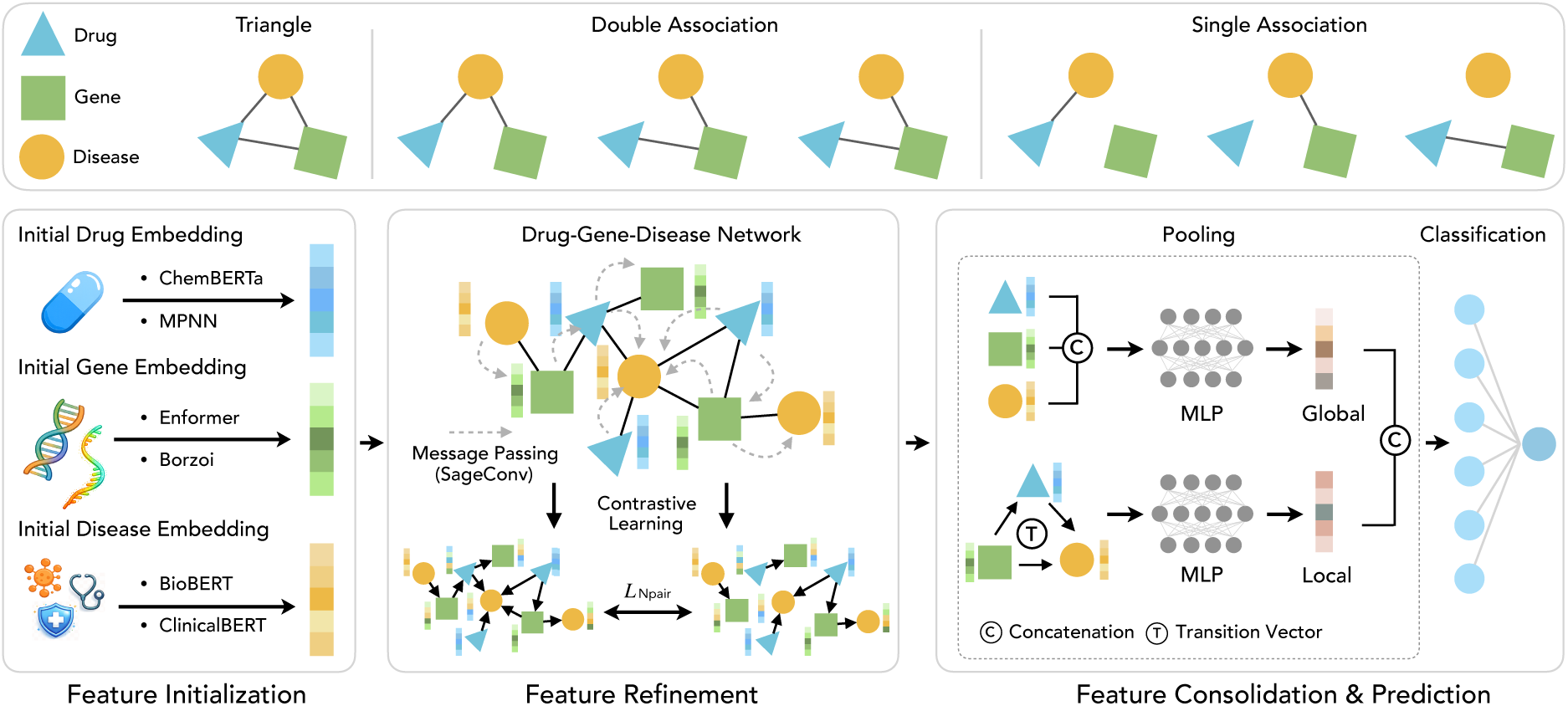
HANAMI workflow. HANAMI is designed to predict seven drug-gene-disease motifs through three stages. Feature initialization: multi-modal biomedical entities are encoded using domain-specific models, including ChemBERTa and MPNN for drugs, Enformer and Borzoi for genomic data, and BioBERT and ClinicalBERT for diseases. Feature refinement: initialized embeddings are refined through a relation-aware drug-gene-disease network using GraphSAGE-based message passing, optimized via an *N*-pair contrastive learning objective (*L*_Npair_). Feature consolidation and prediction: the final representation is synthesized through structure-aware aggregation (global concatenation and local semantic vectors) and passed to an MLP classifier to identify structural motifs such as triangle and star associations.

### Performance comparison and analysis

HANAMI demonstrates robust predictive performance across diverse drug-gene-disease motif types, consistently outperforming TriMoGCL and other baseline methods. On the MS dataset (Fig. 2), HANAMI identifies triangle motifs with a median AUROC of 0.87, compared with 0.81 for TriMoGCL (*P*-value = 3.7E-2). For double-association motifs (disease-star, drug-star, and gene-star), HANAMI reaches a median AUROC of ∼0.95, representing a 3–5% improvement over TriMoGCL (*P*-values = 5.9E-4, 1.4E-5, and 5.7E-3, respectively). Single-association tasks achieve median AUROCs of 0.99 for drug-disease, 0.93 for gene-disease, and 0.98 for gene-drug (*P*-values = 4.0E-2, 2.1E-4, and 2.0E-3, respectively). In addition to higher predictive accuracy, HANAMI also exhibits substantially lower variance across repeated experiments. These advantages persist on the large-scale DRKG benchmark (Fig. 3). HANAMI achieves a median AUROC of 0.93 for triangle motifs versus 0.91 for TriMoGCL (*P*-value =1.3E-5), 0.94–0.96 for double-association motifs compared with 0.91–0.93 (*P*-values = 5.6E-10, 7.0E-9, and 6.2E-10, respectively), and ∼0.98 for single-association (*P*-values = 7.9E-6, 6.0E-10, and 6.3E-8, respectively). Similar gains were observed using AUPRC (Supplementary Figs. 1 and 2), confirming that HANAMI reliably mitigates false-positive discoveries within imbalanced drug-gene-disease motifs. Together, these results indicate that HANAMI effectively integrates global network topology, local relational features, and rich domain knowledge, providing accurate and robust motif-level predictions across complex biomedical networks.

**Fig. 2.**
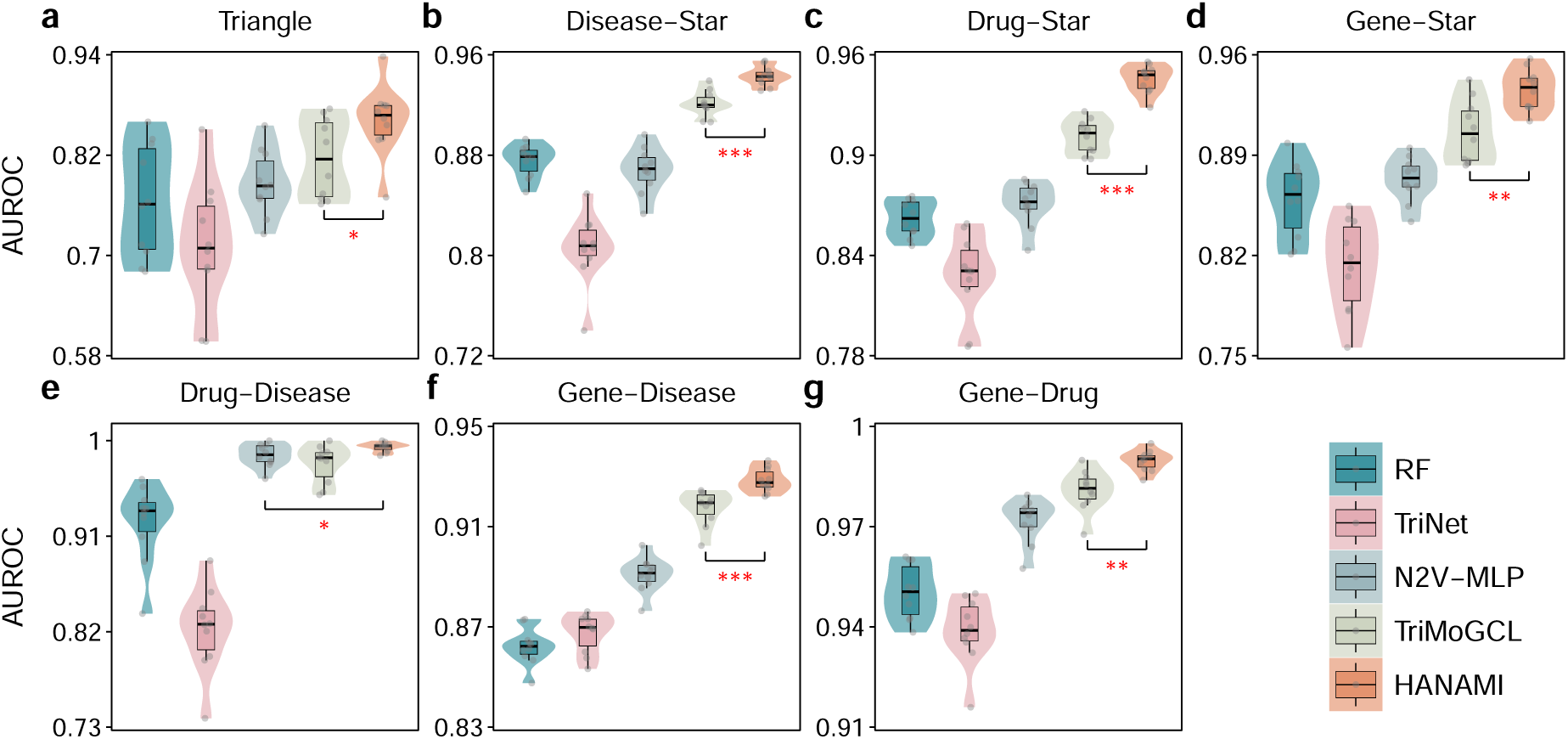
Performance comparison on the MS dataset. **a–g.** AUROC scores for seven structural motifs (triangle, disease-star, drug-star, gene-star, drug-disease, gene-disease, and gene-drug) across RF, TriNet, N2V-MLP, TriMoGCL, and HANAMI on the small-scale MS network. Statistical significance of performance differences is indicated by asterisks (* *P*-value *<* 0.05; ** *P*-value *<* 0.01; *** *P*-value *<* 0.001).

**Fig. 3.**
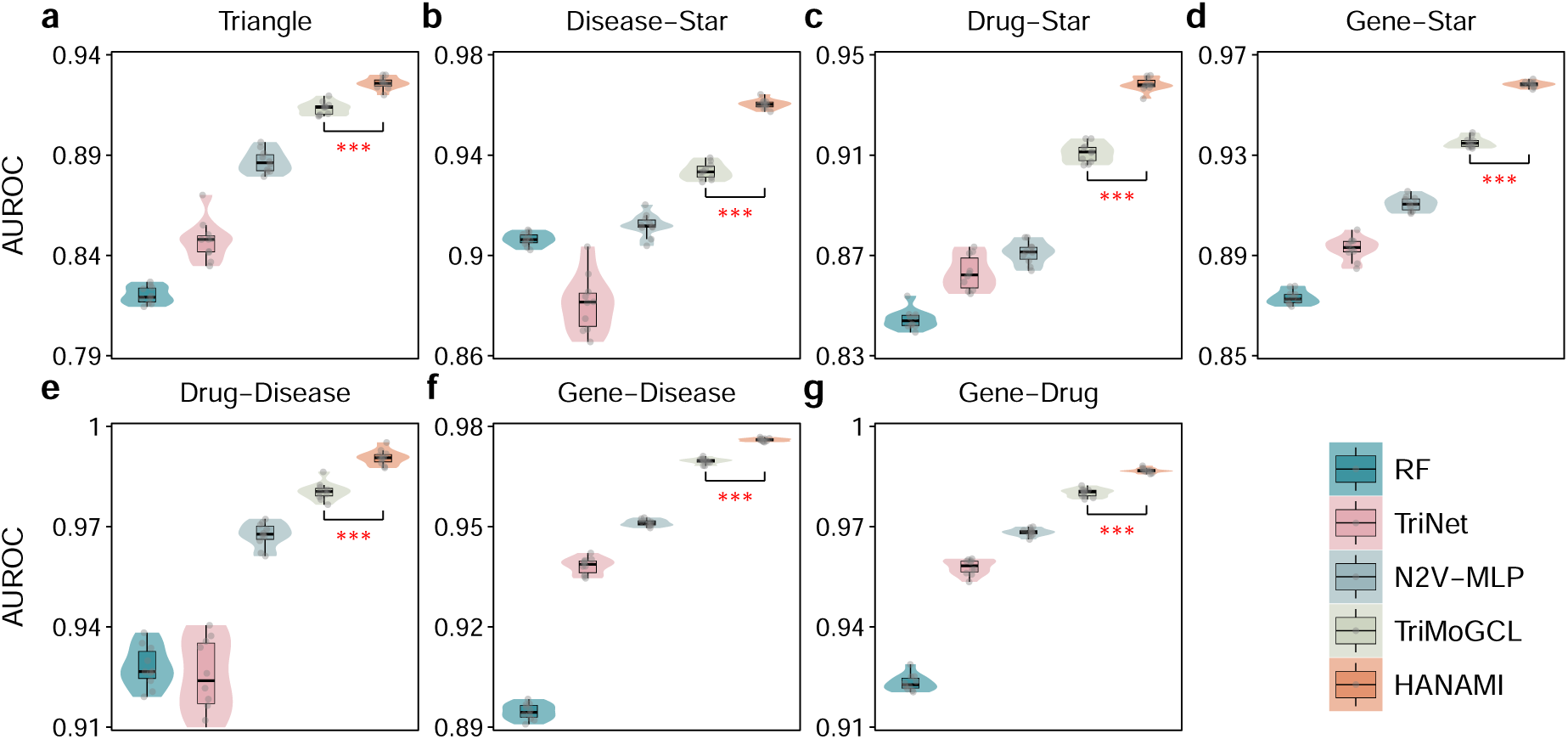
Performance comparison on the DRKG dataset. **a–g.** AUROC scores for seven structural motifs (triangle, disease-star, drug-star, gene-star, drug-disease, gene-disease, and gene-drug) across RF, TriNet, N2V-MLP, TriMoGCL, and HANAMI on the large-scale DRKG network. Statistical significance of performance differences is indicated by asterisks (* *P*-value *<* 0.05; ** *P*-value *<* 0.01; *** *P*-value *<* 0.001).

### Validation with unseen drugs, diseases and genes

We evaluated the inductive generalization capability of HANAMI on entities that were absent from the training data. Under these cold-start conditions, HANAMI significantly outperforms traditional classifiers (Random Forest and N2V-MLP), the multi-branch fusion approach (TriNet), and the advanced graph neural network approach (TriMoGCL) (Fig. 4). In the triangle motif prediction task, HANAMI achieves an AUROC of 0.76, exceeding the second-best method TriMoGCL by 0.16 (*P*-value = 5.4E-4). In double-association prediction tasks, HANAMI attains an average AUROC of 0.86, representing a 0.23 improvement over TriMoGCL (*P*-values = 1.4E-6, 7.4E-10, and 2.1E-7 for disease-star, drug-star, and gene-star, respectively). For single-association configurations, HANAMI maintains an average AUROC of 0.92, outperforming TriMoGCL by 0.13 (*P*-values = 3.8E-4, 6.9E-7, and 2.0E-5 for drug-disease, gene-disease, and gene-drug, respectively). Similar improvements were also observed in AUPRC (Supplementary Fig. 3). These results suggest that HANAMI can effectively generalize structural knowledge learned during training to previously unseen nodes. This capability is critical for practical drug discovery scenarios, where newly developed compounds and understudied disease indications continually emerge.

**Fig. 4.**
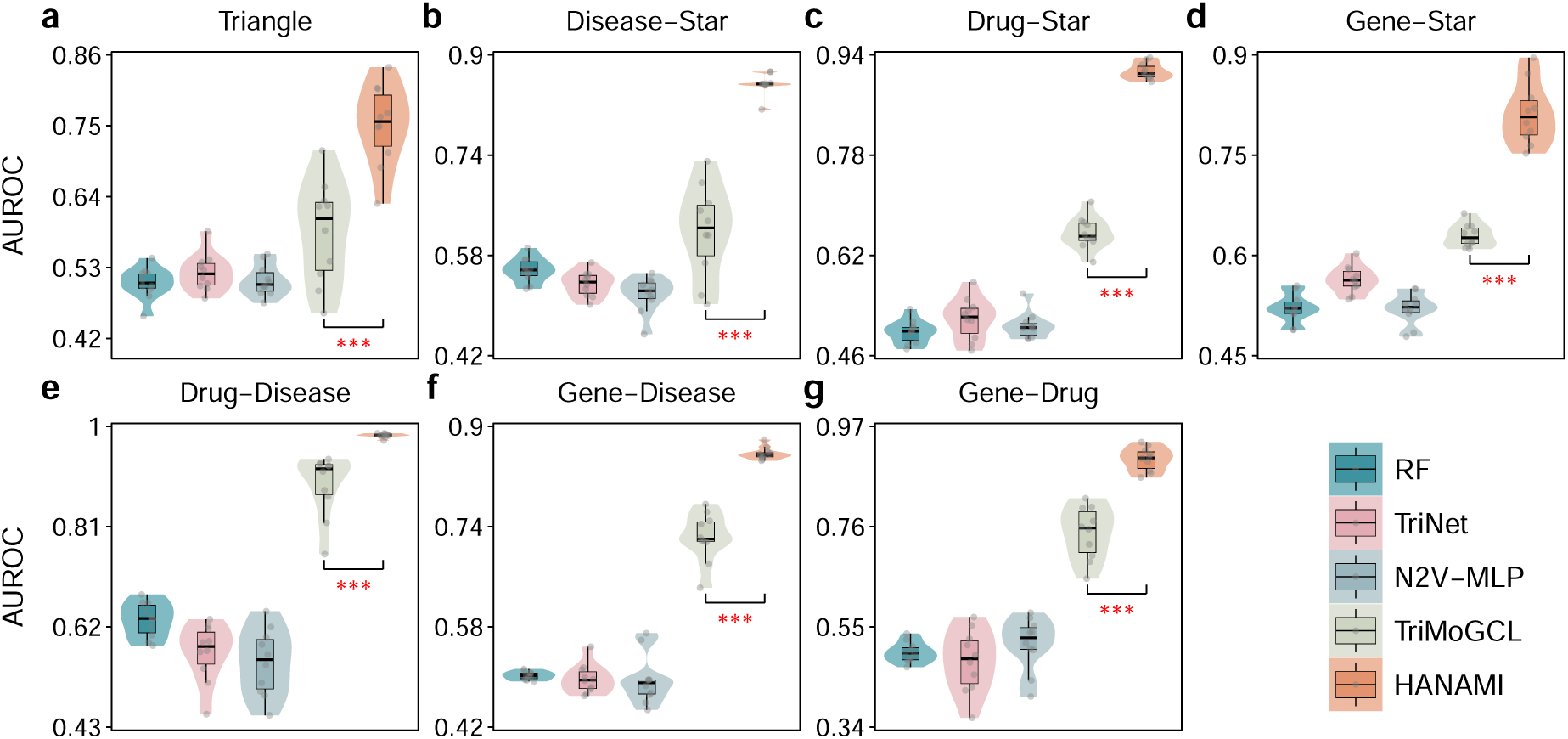
Inductive generalization performance under the cold-start setting. **a–g.** AUROC scores for seven structural motifs (triangle, disease-star, drug-star, gene-star, drug-disease, gene-disease, and gene-drug) across RF, TriNet, N2V-MLP, TriMoGCL, and HANAMI under cold-start conditions. Statistical significance of performance differences is indicated by asterisks (* *P*-value *<* 0.05; ** *P*-value *<* 0.01; *** *P*-value *<* 0.001).

### Biological interpretation and clinical concordance

We evaluated whether HANAMI could prioritize drug and disease relations investigated in clinical studies and provide gene-level biological context through the corresponding motifs. Because confirming candidate relations against individual trial records required extensive record-level review, we focused this analysis on the MS dataset. We compared the ranks assigned by HANAMI and the four baselines for the complete cohort of 1,630 motifs from 785 drug and disease pairs investigated in Phase II or III treatment studies registered in ClinicalTrials.gov. Lower rank percentiles indicate higher predicted priority. In Fig. 5a, HANAMI ranked 524 motifs best among the five methods. These motifs represented 32.15% of the complete cohort, 112 more than TriMoGCL (*P*-value = 1.5E-2). In Fig. 5b, HANAMI achieved a mean rank percentile of 35.69%, which was 0.88 percentage points lower than TriMoGCL (*P*-value = 2.0E-2).

**Fig. 5.**
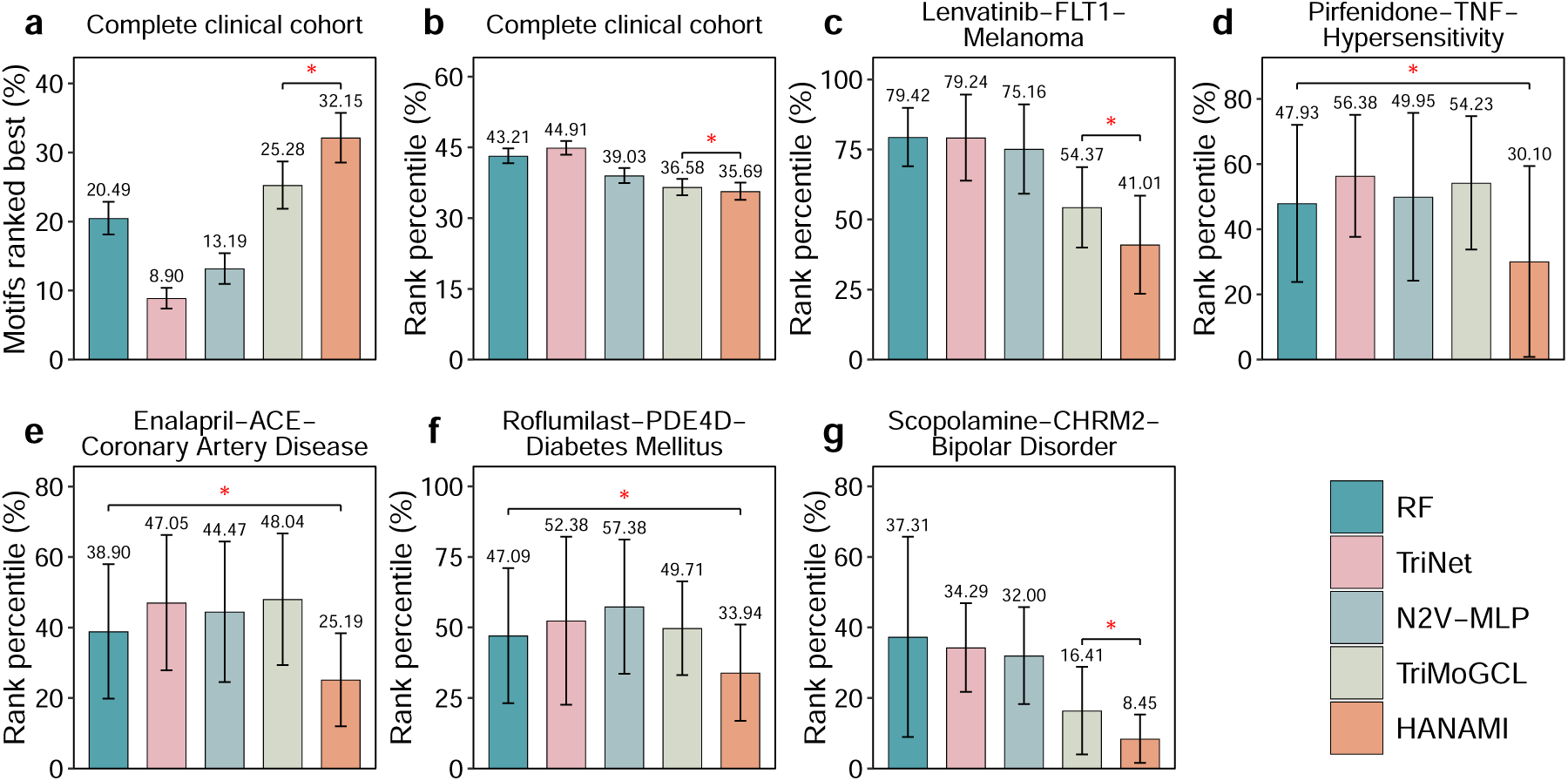
Motif ranking performance on the MS clinical evidence cohort. The complete cohort comprised 1,630 MS gene-star motifs from 785 drug and disease pairs investigated in Phase II or III treatment studies. **a–b.** Percentage of motifs ranked best by each method and mean motif rank percentile across RF, TriNet, N2V-MLP, TriMoGCL, and HANAMI within this cohort. **c–g.** Five representative case studies from different disease types across the same methods. Error bars in a–b show 95% confidence intervals clustered by drug and disease pair, calculated from the ten-seed mean values. In c–g, bars show mean rank percentiles with standard deviations across ten seeds. Exploratory one-sided paired *t*-tests across ten matched seeds compared HANAMI with the baseline having the lowest mean percentile in each panel. Lower percentiles indicate better prioritization. Statistical significance is indicated by red asterisks (* *P*-value *<* 0.05).

To place these cohort-level results in a broader biological context, we grouped the clinically supported pairs and their associated motifs into eight broad disease categories based on the primary disease process and affected organ system. We then selected five representative cases (Fig. 5c–g), prioritizing pathologically complex diseases with high unmet needs, chronic treatment burden, and drug and disease relations investigated in Phase II or III studies registered recently [43–47]. In these illustrative cases, HANAMI assigned the lowest rank percentile among the five methods. The intermediary genes in the motifs revealed two complementary forms of biological context connecting diseases and drugs. Some genes identified direct pharmacological targets, whereas others placed the relations within disease-relevant pathways. The lenvatinib–FLT1–melanoma motif provides the clearest example in which these forms converge. FLT1 (fms related receptor tyrosine kinase 1) encodes VEGFR1 (vascular endothelial growth factor receptor 1), a kinase inhibited by lenvatinib, and experimental melanoma models implicate VEGFR1 in tumor-cell vasculogenic mimicry and growth [48, 49]. By identifying FLT1 rather than only the drug and disease relation, the motif points to a melanoma-cell program beyond the conventional view of endothelial angiogenesis centered on VEGFR2 (vascular endothelial growth factor receptor 2). The other four motifs extend this interpretation across distinct mechanisms. ACE (angiotensin I converting enzyme) links enalapril to renin–angiotensin regulation in coronary artery disease [50], while PDE4D (phosphodiesterase 4D) links roflumilast to cAMP (cyclic adenosine monophosphate) signaling and glucose regulation in diabetes mellitus, where the drug was investigated alongside background diabetes therapy [51, 52]. TNF (tumor necrosis factor) links pirfenidone to inflammatory cytokine regulation, consistent with its suppression of TNF-*α* (tumor necrosis factor alpha) production in experimental models [53]. The trial supporting the MS label Hypersensitivity studied chronic hypersensitivity pneumonitis [54]. CHRM2 (cholinergic receptor muscarinic 2) places scopolamine within muscarinic signaling investigated in bipolar depression. The corresponding trial did not show greater antidepressant benefit than placebo [55]. Together, these cases show how intermediary genes can translate prioritized drug and disease relations into specific, testable biological hypotheses.

## Discussion

In this work, we introduce HANAMI, a multi-view graph neural network framework for predicting heterogeneous motifs among drugs, genes, and diseases. HANAMI incorporates representations from pre-trained language models and domain-specific encoders to capture rich semantic features for each entity type. These features are further refined through GraphSAGE-based message passing and integrated with structure-aware pooling mechanisms that capture relational patterns in the network. In addition, a contrastive learning objective with multiple positive and negative pairs of examples helps the model distinguish subtle structural differences. Together, these components enable HANAMI to uncover complex interaction patterns and produce accurate motif-level predictions in heterogeneous biomedical networks. Benchmark evaluations on the MS and DRKG datasets demonstrate that HANAMI consistently surpasses state-of-the-art methods, particularly in scenarios involving unseen or rare entities. Analysis of the complete clinical evidence cohort extends the benchmark findings by showing that HANAMI can prioritize clinically investigated drug and disease relations more effectively than the strongest baseline and connect them to gene-level hypotheses for biological validation.

To assess how HANAMI’s architectural components contributed to benchmark performance, we conducted a systematic ablation study using the MS dataset. The performance of the model on seven distinct topological motif structures is summarized in Table 3. We benchmarked HANAMI against TriMoGCL using three ablation variants to isolate the effect of each component: (i) HANAMI-w/o-*N*-pair, which replaces the *N*-pair objective with the standard contrastive learning approach used in TriMoGCL; (ii) HANAMI-w/o-GraphSAGE, which replaces the GraphSAGE-based message passing strategy with the regular graph convolution architecture; (iii) HANAMI-w/o-embedding, which removes the comprehensive initial entity embeddings and reverts to the initialization strategy used in TriMoGCL. Among the tested variants, the removal of GraphSAGE-based message passing results in the most pronounced performance decline across all motif structures. The absence of comprehensive initial embeddings also leads to a substantial reduction in AUROC, particularly for the triangle structure. Finally, replacing the *N*-pair objective with standard contrastive learning results in a moderate decrease in performance. Overall, the ablation study indicates that the three tested components contribute differently to predictive accuracy on MS.

HANAMI introduces several architectural improvements that distinguish it from existing methods such as TriMoGCL for biomedical motif prediction. First, HANAMI systematically leverages available pre-trained language models and domain-specific encoders to generate rich, multi-modal embeddings that capture semantic, chemical, and structural context for drugs (ChemBERTa-3, MPNN), genes (Borzoi, Enformer), and diseases (BioBERT, ClinicalBERT). In contrast, TriMoGCL relies on more limited domain knowledge and simpler feature representations, which may not fully capture complex biochemical and functional relationships among biomedical entities. Second, HANAMI employs a GraphSAGE-based architecture that models the distribution of local neighborhoods rather than learning absolute node coordinates. By contrast, TriMoGCL uses traditional Laplacian-based graph convolutions, which are prone to oversmoothing and often struggle in inductive settings. HANAMI’s inductive design not only supports prediction for previously unseen entities but also stabilizes training through weight matrix basis decomposition, while preserving local topological and relational information. Finally, HANAMI optimizes the latent space using an *N*-pair contrastive objective, which simultaneously evaluates multiple negative samples to sharpen decision boundaries and prevents feature collapse in rare motifs. In comparison, TriMoGCL relies on pairwise contrastive learning and can be sensitive to augmentation strategies, potentially producing biased embeddings in less frequent motifs. These predictive gains came with a computational tradeoff. Across the seven motif prediction tasks, HANAMI required more training time than TriMoGCL on MS (16.06 versus 12.96 minutes) and DRKG (265.93 versus 85.66 minutes), but used less peak allocated GPU memory on both datasets (1.19 versus 1.47 GiB and 4.66 versus 7.21 GiB, respectively).

HANAMI offers a powerful computational framework for exploring the intricate interplay among drugs, genes, and diseases, with broad implications for biomedical research. It identifies previously unannotated motifs and generates biological hypotheses about the links among drug targets, genes, and diseases. Moreover, HANAMI can perform zero-shot inference on new or rare entities, allowing the evaluation of therapeutic potential even when prior experimental data are lacking. Beyond the benchmark evaluations, the clinical analysis further suggests that prioritized motifs can guide researchers toward both drug and disease relations worth investigating and specific molecular targets or pathways to examine. By capturing both global network topology and local relational structures, the framework supports systematic prioritization of high-value drug candidates, the repurposing of existing therapeutics, and the identification of drug-gene-disease interactions, offering a rational foundation for experimental validation. Beyond drug discovery, HANAMI can map broader molecular, genomic, and clinical landscapes, inform mechanistic hypotheses, and guide precision medicine initiatives, as well as translational research efforts aimed at understanding complex disease processes.

Despite these strengths, HANAMI has several limitations that warrant future investigation. First, the framework relies on existing biomedical databases and curated annotations, which may be incomplete or biased. Integrating additional data sources, such as emerging high-throughput multi-omics datasets (e.g., single-cell data and microbiome data) and electronic health records from patients, could further enhance its coverage and predictive performance. Second, HANAMI currently treats interactions as static snapshots, whereas many biological processes are dynamic and context-dependent. Extending the model to incorporate temporal or condition-specific networks would allow for a more realistic representation of disease progression and drug response. Third, while the framework captures local and global network dependencies, it does not explicitly model higher-order biochemical constraints, such as metabolic flux or protein complex stoichiometry. Integrating mechanistic priors or physics-informed constraints could enhance interpretability and biological relevance. Fourth, although the model demonstrates robust performance in zero-shot scenarios, systematic literature-based and experimental validation of its predictions is needed to confirm their translational relevance. Finally, the clinical evidence analysis was limited to the small network, so its generalizability to other biomedical networks remains to be assessed. The biological hypotheses associated with the shared genes also require experimental validation. Overall, HANAMI establishes a compelling methodology for drug-gene-disease motif prediction, clinically grounded motif prioritization, and biological interpretation by bridging molecular semantics with higher-order network topology, offering a robust foundation for therapeutic target identification and the study of complex biomedical interactions.

## Methods

### HANAMI model

#### Feature initialization

HANAMI relies on translating multi-modal biomedical data into a unified and computationally efficient representation to perform relational reasoning. To achieve this, the framework encodes the intrinsic properties of drugs, genes, and diseases into a shared high-dimensional embedding space using deep learning architectures tailored to specific data types, including chemical SMILES (simplified molecular input line entry system) strings [56], genomic sequences, and clinical terminology. This strategy preserves critical biological information, such as molecular topology, genomic regulatory signals, and disease phenotypes, as dense feature vectors.

##### Drug initial embeddings

For small molecules, embeddings are extracted from SMILES strings to represent molecular architecture as text-based sequences. The framework utilizes ChemBERTa-3 [35], a biochemical language model, to capture chemical semantics, producing hidden states that are aggregated into a 1152-dimensional vector. In parallel, we apply an MPNN [36] to capture structural information from molecular graphs. In the setting, SMILES strings are converted into graph representations, where atom states are iteratively updated through vector exchanges with neighboring nodes to capture local chemical environments and atomic connectivity. This graph-based procedure produces a 300-dimensional vector representation for each drug. The final step involves the concatenation of the two feature vectors, resulting in a comprehensive 1452-dimensional embedding for each drug node. By integrating semantic information from molecular sequences with topological information from molecular graphs, this representation provides a rich and complementary initial feature state for drug nodes.

##### Gene initial embeddings

Gene features are generated by querying the NCBI (National Center for Biotechnology Information) [57] database for chromatin coordinates to retrieve DNA sequences from the reference genome. These sequences are standardized to 524,288 base pairs for Borzoi [38] and 114,688 for Enformer [37] to predict genomic tracks of 4,096-by-7,611 and 896-by-5,313 dimensions, respectively. To manage this high dimensionality, principal component analysis [58] is used to reduce the feature space to 100 components per track while retaining significant variance. These compressed matrices are processed by a convolutional neural network-based autoencoder [59] to extract latent representations, yielding a fixed 1024-dimensional vector for each gene. Finally, the latent vectors from both models are fused via concatenation and projected through an MLP to create a unified 1024-dimensional embedding that integrates complementary regulatory signals captured by both sequence models.

##### Disease initial embeddings

Disease representations are constructed by combining clinical text features with structured categorical data. To extract features from disease names and clinical narratives, biomedical language models BioBERT [39] and ClinicalBERT [40] are used to generate 1024-dimensional and 768-dimensional embeddings, respectively. These textual features are supplemented by medical subject headings identifiers[60] to integrate standardized medical classification systems. Following established fusion protocols, the language model outputs are concatenated with the medical subject headings identifiers to produce a final 1792-dimensional embedding for every disease node. This approach captures both unstructured medical descriptions and structured hierarchical categories within the initial feature space.

##### Input dimension alignment

The resulting drug, gene, and disease embeddings have dimensions of 1,452, 1,024, and 1,792, respectively. To give all node types the same input dimension, the drug and gene vectors are zero-padded to 1,792 dimensions.

#### Feature refinement

HANAMI refines initial embeddings of drugs, genes, and diseases through a heterogeneous graph contrastive learning framework. The combined feature matrix is batch normalized [61] to standardize each feature dimension across nodes before the first GraphSAGE [41] layer maps all node types to a shared hidden dimension. We then employ a relation-aware topology encoder rooted in the GraphSAGE framework to refine node representations. The architecture consists of two stacked layers with distinct aggregation mechanisms to capture multiscale structural information. In contrast to spectral graph convolution networks that depend on the full graph Laplacian, GraphSAGE functions inductively by aggregating information from a node’s local neighborhood, which naturally accommodates heterogeneous multimodal features and generalizes to unseen entities. Information is aggregated from the complete neighborhood of each node rather than a fixed-size sampled subset, preserving the complete local topology required for motif prediction while avoiding the additional variance and hyperparameter introduced by neighborhood sampling. For any node *v* at layer *l*, we aggregate neighbor representations via

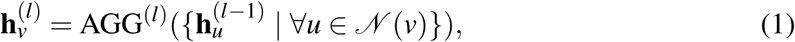

where 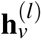 denotes the embedding of node *v* at layer *l*, AGG^(*l*)^(·) denotes the aggregation function at layer *l* (such as maximum pooling or mean pooling), and *N* (*v*) represents the neighbor set of node *v*. This aggregated neighborhood context is then concatenated with the node’s existing state to update its embedding

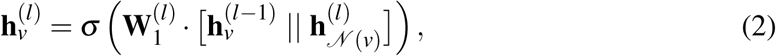

where 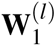 denotes the learnable weight matrix at layer *l*, the operator “||” represents concatenation, and *σ* denotes a nonlinear activation function. The initial embedding 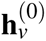 is initialized using the modality-specific feature vector obtained during the feature initialization phase. To ensure parameter efficiency without sacrificing expressiveness, we factorize the weight matrix using basis decomposition 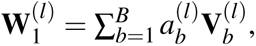, where 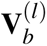 are shared basis matrices and 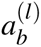 are learnable coefficients. After several layers, the architecture produces a final node embedding vector **h***_v_* that captures both intrinsic semantics and multiscale topological context propagated through the heterogeneous graph.

To regularize the latent space and optimize structural motif representations, we implement an *N*-pair[42] contrastive learning objective. An augmented graph is generated by randomly dropping edges, producing alternative views of the triplets. The contrastive loss seeks to maximize the similarity between positive pairs while minimizing similarity with negative pairs

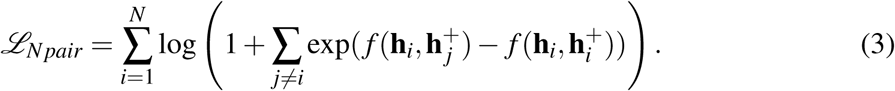

This objective will be integrated into the motif prediction stage to enhance discriminative power. By bringing embeddings of the same triplet closer while distancing unrelated triplets, *N*-pair contrastive learning sharpens the separation between motif classes in the latent space. Consequently, HANAMI becomes more robust to network noise, less prone to overfitting, and better able to detect subtle structural differences.

#### Feature consolidation & prediction

HANAMI constructs comprehensive triplet descriptors by performing an entity-wise aggregation across the refined feature space, thereby synthesizing the specific structural semantics required for classification. First, a global aggregation mechanism effectively fuses the refined embeddings of the distinct nodes forming a triplet into a unified context vector. For a triplet *t* comprising a disease node *i*, a drug node *j*, and a gene node *k*, their hidden embeddings are concatenated as **x***_t_* = [**h***_i_* || **h** *_j_* || **h***_k_*]. This combined vector is subsequently transformed by an MLP to generate the final global-consolidated representation, denoted by **x̃***_t_*. To explicitly capture local interaction features, we further utilize a local aggregation mechanism inspired by triangle Lasso[62]. Specifically, semantic vectors are constructed to characterize the relational dynamics between node pairs. These pairwise relations are mapped into a latent space through a nonlinear transformation and then concatenated as

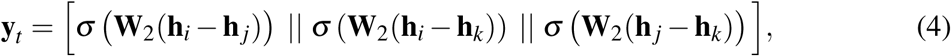

where **W**_2_ denotes the learnable weight matrix. The resulting local interaction representation is further refined through an MLP to yield **ỹ***_t_*. The macro and micro topological aggregation mechanisms produce complementary triplet-level representations, capturing both node-specific context and inter-node relational dynamics.

The final triplet embedding **z***_t_*is synthesized by concatenating the global-consolidated embedding **x̃***_t_*and the local-consolidated embedding **ỹ***_t_* into a unified vector **z***_t_* = [**x̃***_t_* || **ỹ***_t_*]. To determine the motif category, a linear transformation projects these features into the label space, followed by softmax normalization to compute the confidence score

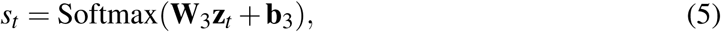

where **W**_3_ denotes the learnable weight matrix linking the feature space to the binary output classes, and **b**_3_ is the corresponding bias term. The architecture is trained by minimizing a composite loss function

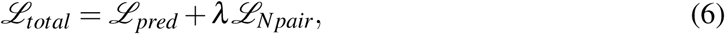

where *L_pred_* and *L_N pair_* denote the binary cross-entropy loss and the auxiliary *N*-pair contrastive loss, respectively, and *λ* ∈ [0, 1] is a tuneable parameter. This objective configuration ensures the model retains semantic motifs while filtering network noise, providing stable predictions for diverse biological configurations.

### Experimental details

#### Datasets

To evaluate the performance of HANAMI, two complementary biomedical knowledge graphs were employed: the multi-scale interactome (MS) [33] and drug repurposing knowledge graph (DRKG) [34] datasets. The two datasets were selected to capture pharmacological associations at both systemic macroscopic level and granular molecular level. The MS dataset consists of 29,959 nodes organized into four categories (drugs, proteins, diseases, and Gene Ontology biological functions) and contains 478,728 edges across four relation types (i.e., drug-protein, disease-protein, protein-protein, and protein-biological function), while DRKG contains 97,238 biological entities spanning 13 distinct types and forms a densely connected network with 5,874,261 triplets across 107 relation types. To ensure precise modeling of structural interactions, specific subgraphs composed exclusively of edges connecting drug, gene (or protein), and disease entities were extracted from these original networks. Quantitative characteristics of the resulting graph structures derived from the MS and DRKG datasets are summarized in Table 1, including the counts of each node type (drugs, genes, and diseases), the total number of edges, and the frequencies of structural motifs.

**Table 1:**
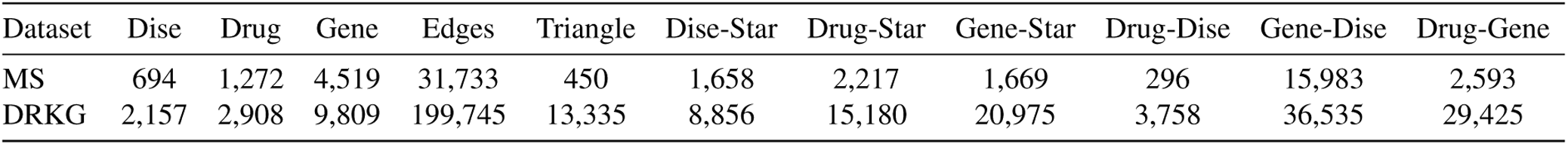
The statistics of the MS and DRKG datasets in one triplet sampling.

| Dataset | Dise | Drug | Gene | Edges | Triangle | Dise-Star | Drug-Star | Gene-Star | Drug-Dise | Gene-Dise | Drug-Gene |
| --- | --- | --- | --- | --- | --- | --- | --- | --- | --- | --- | --- |
| MS | 694 | 1,272 | 4,519 | 31,733 | 450 | 1,658 | 2,217 | 1,669 | 296 | 15,983 | 2,593 |
| DRKG | 2,157 | 2,908 | 9,809 | 199,745 | 13,335 | 8,856 | 15,180 | 20,975 | 3,758 | 36,535 | 29,425 |

We followed the motif sampling procedure described in [30]. Given that a single edge can theoretically participate in multiple triplet motifs, leading to an exponential explosion of data, a simplified sampling rule was adopted to assign each edge to a maximum of one triplet. To reduce potential bias introduced by this simplification and ensure evaluation integrity, 10 independent triplet sampling folders were conducted for each motif, and the final results were then averaged to evaluate model robustness. The sampling procedure prioritized complex structural motifs to maintain dense topological characteristics, specifically targeting triangles, disease-stars, drug-stars, and gene-stars. Following this priority order, triplets within each category were iterated through in a randomized sequence. If all individual edges within a specific triplet had not been previously visited, the triplet was collected for evaluation. After exploring complex motifs, any remaining unassigned edges were allocated to disconnected nodes to form single-edge motifs. For example, an unassigned drug-disease edge could be combined with an unconnected, randomly sampled gene node to construct a drug-disease motif, ensuring the entire network is utilized without redundant edge participation.

#### Baselines

To evaluate the performance of HANAMI, we compared it against a set of baseline methods, including traditional and state-of-the-art methods. These baseline methods include traditional ensemble learning, random walk-based hybrid models, multi-branch fusion architectures, and advanced graph neural networks:

- TriMoGCL [30] is a state-of-the-art graph neural network method designed for predicting triplet motifs in disease-drug-gene interactions. The framework combines graph convolutional encoders with contrastive learning and aggregation mechanisms to effectively capture structural patterns within biomedical networks.
- TriNet [29] is a multi-branch fusion framework designed for modeling triplet interactions in biomedical networks. The method utilizes parallel network branches to transform node embeddings before synthesizing them via concatenation and nonlinear projection layers to capture complex structural dependencies.
- N2V-MLP [28] is a hybrid baseline that exploits the Node2Vec algorithm to extract node features by capturing long-distance topological dependencies via random walks, which are subsequently fed into an MLP for motif classification.
- Random forest (RF) [19] is a traditional ensemble method used as a baseline. Each triplet is represented by a feature vector combining raw node attributes, pairwise absolute differences to capture interactions, and structural metrics such as common neighbors and Jaccard coefficients for intra-triplet node pairs.

#### Evaluation protocol and cold-start design

Data partitioning into training, validation, and testing sets was tailored to the scale and complexity of each biological network. For the MS dataset, triplets for each motif were split using an 8:1:1 ratio, while the larger DRKG dataset employed a 4:3:3 split to prevent metric ceiling effects and ensure a more rigorous evaluation of the model’s generalization capabilities. For binary classification, negative samples for each target motif were generated from equal proportions of triplets in non-target motif categories, forcing the model to learn motif-specific structural signatures rather than simple connectivity. These partitioning and negative sampling strategies allow evaluation across both local physical interactions and global associations, demonstrating generalizability across different drug repurposing contexts and network distributions. To ensure unbiased assessment, stratified cross-validation was applied. Each fold was sequestered for testing while the remaining data trained the model, iterating until every association was evaluated. For each iteration, the training graph was reconstructed using only edges assigned to the training set, preventing contamination from test entities during feature refinement. Additionally, the model was evaluated against the full set of non-validated pairs rather than balanced sub-sampling, more accurately reflecting the challenge of detecting rare therapeutic signals in large-scale interaction spaces.

Predicting therapeutic indications for new compounds remains a major challenge for graph-based models, commonly referred to as the strict cold-start problem. To address this, we incorporated an inductive transfer learning strategy [63], which leverages feature similarities within pre-trained embeddings to extend the learned relational patterns to novel nodes. In this setting, the training and validating datasets were established using a subgraph of DRKG. This subgraph was constructed by excluding all nodes present in MS, and all edges connected to these entities were removed to prevent data leakage. By strictly isolating the training and validating data, the MS dataset served as a completely independent test set for evaluating performance on the target domain. During inference, unseen entities were projected into a shared latent space through their respective encoders, enabling prediction without model re-training.

#### Biological analysis details

We analyzed gene star candidates in MS that connected a drug and disease through a shared gene but lacked a direct relation between the drug and disease. Within this candidate pool, a pair qualified when a Phase II or III treatment study registered in ClinicalTrials.gov [64] listed the drug as an intervention and the disease as a condition and identified the drug as a treatment for that disease in the trial title. Pairs supported only by uses unrelated to the disease indication, such as comorbidity treatment, supportive care, or toxicity management, were therefore excluded. All qualifying pairs were retained for the primary analysis. The complete cohort therefore contained 1,630 motifs representing 785 drug and disease pairs across 109 disease labels.

Within this cohort, we retained every corresponding motif for pairs with multiple shared genes and evaluated the same motifs with all five methods. For each of ten seeds, each method ranked the same 46,704 MS gene star candidates from highest to lowest prediction score. The score was the triangle classifier output, with higher values indicating stronger support for the triangle class. Rank percentile was calculated as 100 times the descending rank divided by 46,704, with tied scores assigned their average rank. The mean rank percentile across seeds was calculated, and for each motif, the method with the lowest mean rank percentile was counted as the best-ranked method. For biological interpretation, the 109 disease labels were grouped into eight broad categories comprising infectious diseases, neoplastic diseases, cardiovascular diseases, metabolic and endocrine diseases, immune-mediated and inflammatory diseases, neurological and psychiatric diseases, respiratory and gastrointestinal diseases, and genetic, congenital and rare diseases. Each label was assigned according to its primary disease process and affected organ system. Five clinically documented motifs from different disease types were selected as case studies, prioritizing pathologically complex diseases with high unmet needs and chronic treatment burden, to illustrate how the shared gene linked the drug and disease.

#### Evaluation metrics

We evaluated model performance using two statistical measures: the area under the receiver operating characteristic curve (AUROC) and the area under the precision-recall curve (AUPRC). While AUROC provides a general assessment of the model’s ability to rank true associations above random pairs, AUPRC offers a complementary evaluation of performance in the context of highly imbalanced datasets, where positive signals are significantly outnumbered by unknown pairs. These metrics allow us to determine the model’s precision and its resilience against false-positive discoveries. Statistical significance was systematically determined using a one-tailed paired *t*-test. For the clinical evidence analysis, we evaluated model performance using two ranking measures: mean motif rank percentile and the percentage of motifs ranked best by each method. Lower mean percentiles indicated stronger overall prioritization, whereas a higher percentage of motifs ranked best indicated more frequent top performance. HANAMI and TriMoGCL were compared using paired differences in per-motif indicators of being ranked best for Fig. 5a and mean rank percentiles for Fig. 5b. Both comparisons used one-sided *t*-tests with CR1 standard errors clustered by drug and disease pair [65].

#### Hyperparameters

Hyperparameters for HANAMI were determined through an extensive grid search on a dedicated validation set. The search explored multiple embedding dimensions for drugs, diseases, and genes, as well as variations in GraphSAGE layers, aggregation strategies, and unit sizes across both global and local components. Regularization was tuned via standard and attention-specific dropout, while optimization employed the Adam optimizer with a range of learning rates and weight decay values. These choices were made to maximize predictive stability and ensure robust, discriminative representation learning across diverse motif types. Detailed hyperparameter search space and the selected values are provided in Table 2.

**Table 2:**
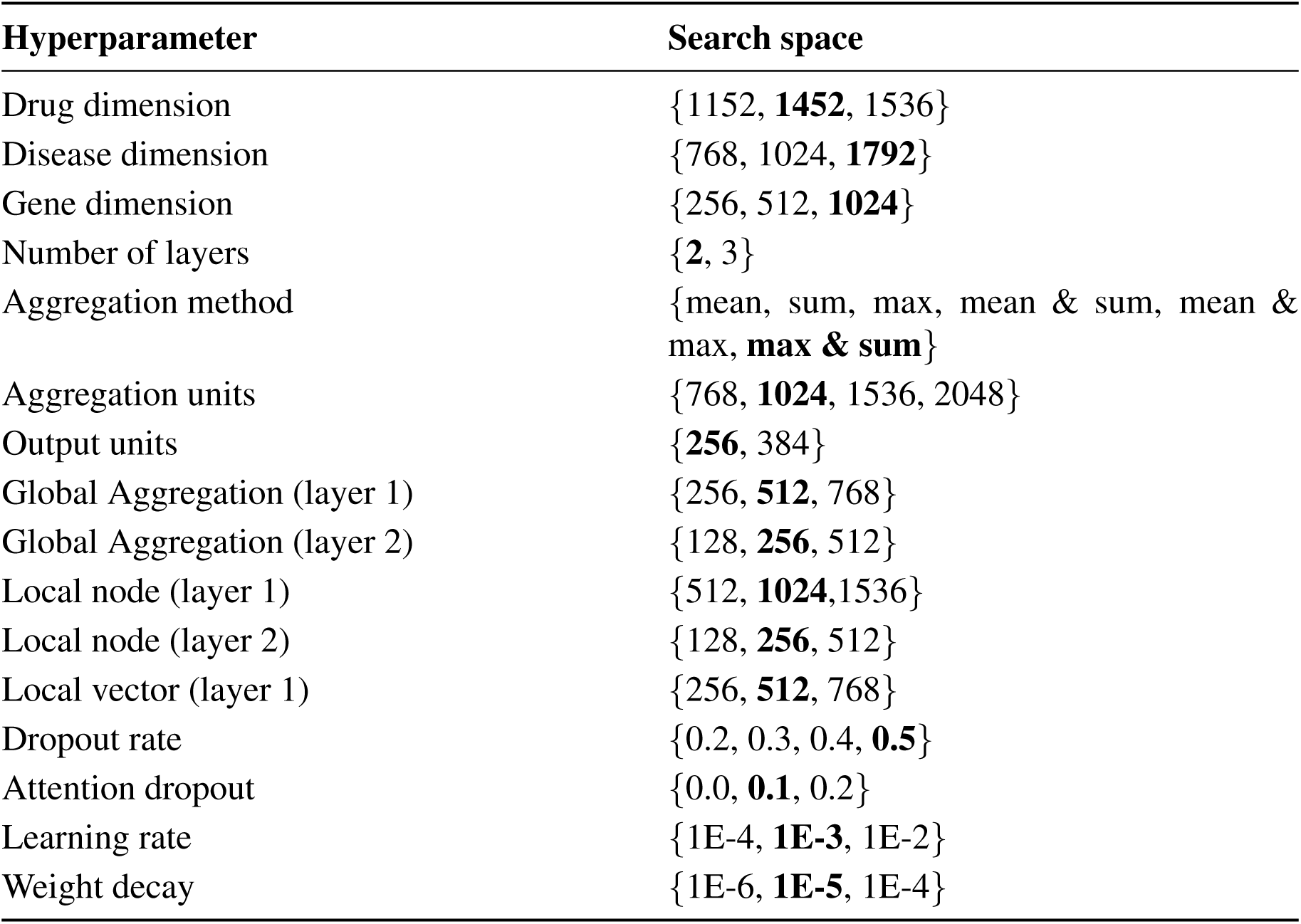
Hyperparameter selection for the proposed framework: Grid search is performed on the validation set, with optimal values shown in bold.

| Hyperparameter | Search space |
| --- | --- |
| Drug dimension | {1152, **1452**, 1536} |
| Disease dimension | {768, 1024, **1792**} |
| Gene dimension | {256, 512, **1024**} |
| Number of layers | {**2**, 3} |
| Aggregation method | {mean, sum, max, mean & sum, mean & max, **max & sum**} |
| Aggregation units | {768, **1024**, 1536, 2048} |
| Output units | {**256**, 384} |
| Global Aggregation (layer 1) | {256, **512**, 768} |
| Global Aggregation (layer 2) | {128, **256**, 512} |
| Local node (layer 1) | {512, **1024**, 1536} |
| Local node (layer 2) | {128, **256**, 512} |
| Local vector (layer 1) | {256, **512**, 768} |
| Dropout rate | {0.2, 0.3, 0.4, **0.5**} |
| Attention dropout | {0.0, **0.1**, 0.2} |
| Learning rate | {1E-4, **1E-3**, 1E-2} |
| Weight decay | {1E-6, **1E-5**, 1E-4} |

**Table 3:**
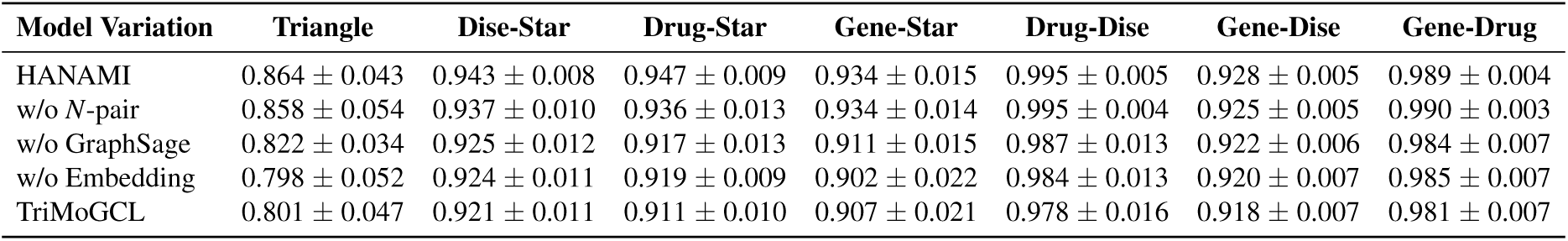
Ablation study results for HANAMI: Average test AUROC are reported with standard deviation over multiple folders.

| Model Variation | Triangle | Dise-Star | Drug-Star | Gene-Star | Drug-Dise | Gene-Dise | Gene-Drug |
| --- | --- | --- | --- | --- | --- | --- | --- |
| HANAMI | 0.864 ± 0.043 | 0.943 ± 0.008 | 0.947 ± 0.009 | 0.934 ± 0.015 | 0.995 ± 0.005 | 0.928 ± 0.005 | 0.989 ± 0.004 |
| w/o *N*-pair | 0.858 ± 0.054 | 0.937 ± 0.010 | 0.936 ± 0.013 | 0.934 ± 0.014 | 0.995 ± 0.004 | 0.925 ± 0.005 | 0.990 ± 0.003 |
| w/o GraphSage | 0.822 ± 0.034 | 0.925 ± 0.012 | 0.917 ± 0.013 | 0.911 ± 0.015 | 0.987 ± 0.013 | 0.922 ± 0.006 | 0.984 ± 0.007 |
| w/o Embedding | 0.798 ± 0.052 | 0.924 ± 0.011 | 0.919 ± 0.009 | 0.902 ± 0.022 | 0.984 ± 0.013 | 0.920 ± 0.007 | 0.985 ± 0.007 |
| TriMoGCL | 0.801 ± 0.047 | 0.921 ± 0.011 | 0.911 ± 0.010 | 0.907 ± 0.021 | 0.978 ± 0.016 | 0.918 ± 0.007 | 0.981 ± 0.007 |

#### Computational cost assessment

We measured training time, inference time, and peak memory for the complete set of seven motif prediction tasks on each dataset, using identical sample splits across methods. Methods ran sequentially on an Intel Core i7-14650HX computer with 32 GB RAM and an RTX 4060 Laptop GPU. Each method retained its original features and graph processing. Neural models trained for 150 epochs per task on the GPU, whereas RF fitted 100 trees on the CPU. Training time included Node2Vec training but excluded data preparation, evaluation, and pretrained feature generation. Inference time was averaged over 20 repetitions after one warm-up, with GPU operations synchronized for timing. Within each dataset, times were summed across tasks, and peak process RAM and allocated GPU memory were reported as maxima across tasks. Computational costs for all five methods are reported in Supplementary Tables 1 and 2.

## Data availability

The processed data and experimental outputs used to support the findings of this study, together with instructions for accessing the original MS and DRKG datasets, are available at: https://github.com/Hongsheng-Xie/HANAMI.

## Code availability

The source code used to implement HANAMI and reproduce the reported analyses is publicly available at: https://github.com/Hongsheng-Xie/HANAMI.

## Author contributions

C.C. and S.K. conceptualized and designed the project and supervised the research. H.X. performed the formal analysis and developed the software. H.X. and Y.G. conducted the investigation and wrote the original draft. S.K. and C.C. reviewed and edited the manuscript. All authors approved the final manuscript for submission.

