## Supplementary information for "Heterogeneous Graph Contrastive Learning for Drug-Gene-Disease Motif Prediction"

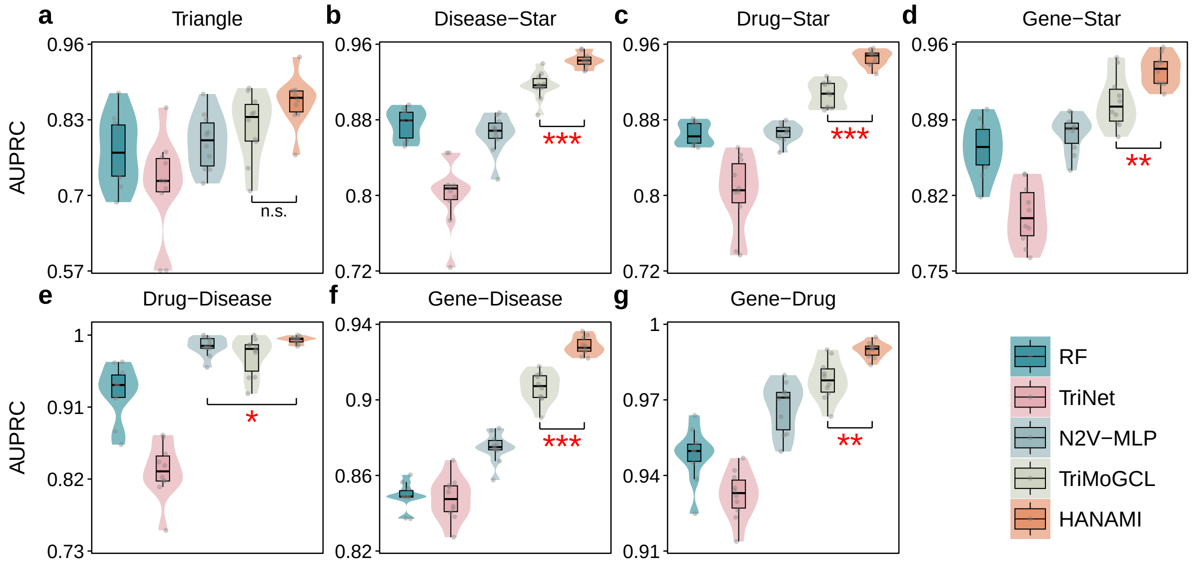


**Supplementary Fig. 1:** Performance comparison on the MS dataset. **a–g.** AUPRC scores for seven structural motifs (triangle, disease-star, drug-star, gene-star, drug-disease, gene-disease, and gene-drug) across RF, TriNet, N2V-MLP, TriMoGCL, and HANAMI on the small-scale MS network. Statistical significance of performance differences is indicated by asterisks (* $P$-value $< 0.05$; ** $P$-value $< 0.01$; *** $P$-value $< 0.001$).

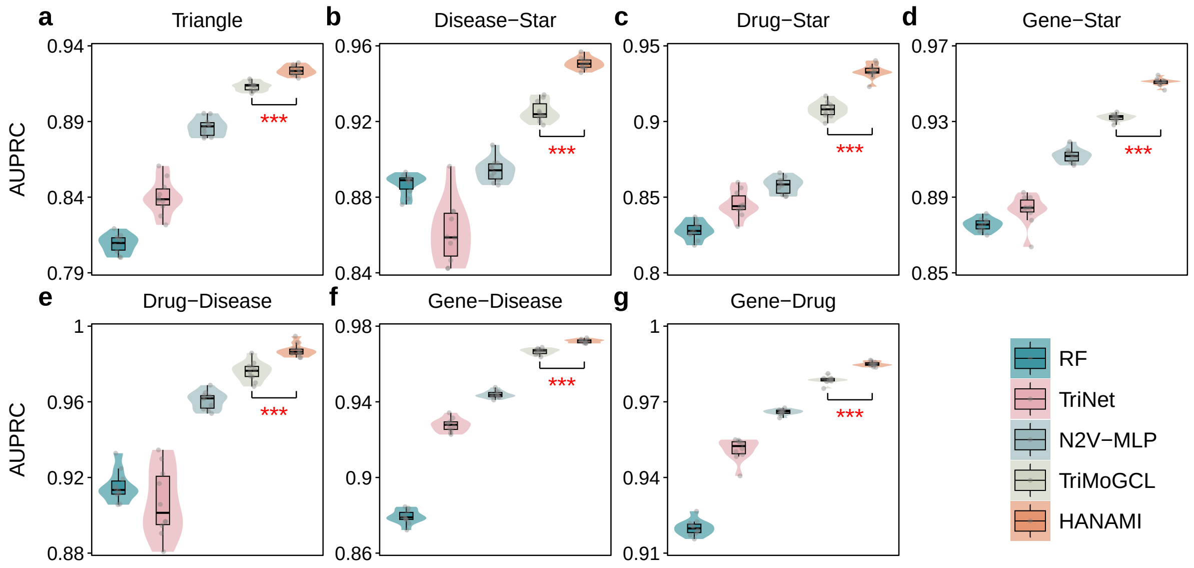


**Supplementary Fig. 2:** Performance comparison on the DRKG dataset. **a–g.** AUPRC scores for seven structural motifs (triangle, disease-star, drug-star, gene-star, drug-disease, gene-disease, and gene-drug) across RF, TriNet, N2V-MLP, TriMoGCL, and HANAMI on the large-scale DRKG network. Statistical significance of performance differences is indicated by asterisks (* $P$-value $< 0.05$; ** $P$-value $< 0.01$; *** $P$-value $< 0.001$).

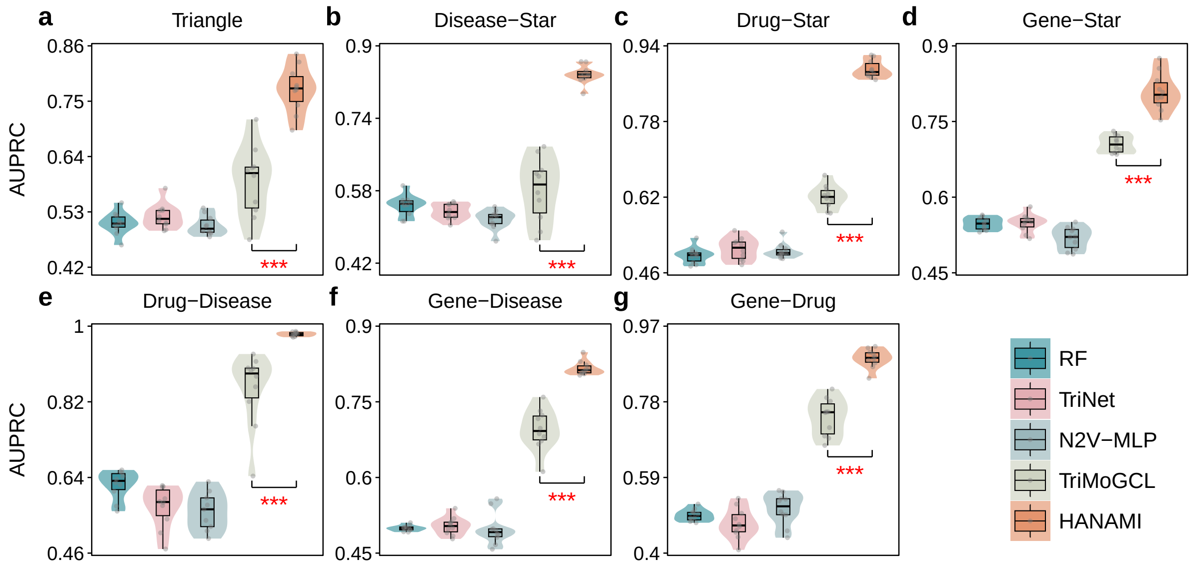


**Supplementary Fig. 3:** Inductive generalization performance under the cold-start setting. **a–g.** AUPRC scores for seven structural motifs (triangle, disease-star, drug-star, gene-star, drug-disease, gene-disease, and gene-drug) across RF, TriNet, N2V-MLP, TriMoGCL, and HANAMI under cold-start conditions. Statistical significance of performance differences is indicated by asterisks (* $P$-value $< 0.05$; ** $P$-value $< 0.01$; *** $P$-value $< 0.001$).

| Method | Training (min) | Inference (ms) | Peak RAM (GiB) | Peak GPU (GiB) |
|---|---|---|---|---|
| RF | 4.29 | 240.49 | 3.84 | — |
| TriNet | 0.45 | 5.79 | 1.29 | 0.37 |
| N2V-MLP | 3.74 | 14.76 | 1.68 | 2.53 |
| TriMoGCL | 12.96 | 57.14 | 1.66 | 1.47 |
| HANAMI | 16.06 | 77.30 | 1.70 | 1.19 |

**Supplementary Table 1:** Computational costs of the five methods on MS. Training and inference times are totals across all seven motif prediction tasks. Memory values represent the maximum across tasks. GPU memory refers to peak allocated memory. RF used the CPU.

| Method | Training (min) | Inference (ms) | Peak RAM (GiB) | Peak GPU (GiB) |
|---|---|---|---|---|
| RF | 570.38 | 1766.22 | 15.90 | — |
| TriNet | 0.82 | 16.68 | 1.98 | 1.81 |
| N2V-MLP | 9.40 | 49.06 | 2.39 | 3.56 |
| TriMoGCL | 85.66 | 157.80 | 3.59 | 7.21 |
| HANAMI | 265.93 | 389.18 | 4.69 | 4.66 |

**Supplementary Table 2:** Computational costs of the five methods on DRKG. Training and inference times are totals across all seven motif prediction tasks. Memory values represent the maximum across tasks. GPU memory refers to peak allocated memory. RF used the CPU.
